# Structural and biochemical characterization of a novel inhibitor of NMNAT1, the gatekeeper of nuclear NAD^+^ biosynthesis

**DOI:** 10.64898/2026.04.07.716846

**Authors:** Carisse Lansiquot, Ruoxi Wu, Joanna P. Davies, Xiangyang Song, H Ümit Kaniskan, Jian Jin, Michael B. Lazarus

**Affiliations:** Department of Pharmacological Sciences, Icahn School of Medicine at Mount Sinai, New York, New York 10029; Mount Sinai Center for Therapeutics Discovery, Icahn School of Medicine at Mount Sinai, New York, New York 10029; Departments of Oncological Science and Neuroscience, The Mount Sinai Tisch Cancer Center, Icahn School of Medicine at Mount Sinai, New York, NY 10029

## Abstract

Nicotinamide adenine dinucleotide (NAD^+^) is crucial for cellular functions including DNA repair and metabolism. Nicotinamide mononucleotide adenylyltransferase (NMNAT) enzymes catalyze the final step of NAD^+^ synthesis from NMN and ATP. There are three NMNAT isoforms: NMNAT1, NMNAT2, and NMNAT3, located in the nucleus, cytoplasm, and mitochondria, respectively. Nuclear NAD^+^ promotes disease progression in NAD^+^-dependent cancers, and it is hypothesized that targeting NMNAT1 with small-molecule inhibitors could be an effective therapeutic strategy. Here, we identify an NMNAT1 inhibitor from a bioactive compound screen and report its effects on NAD^+^ levels and the viability of NMNAT1-dependent cancer cell lines. The compound AMI-1 is a known inhibitor of Protein Arginine N-Methyltransferase 1, and we find that it also inhibits NMNAT1 with similar potency. Additionally, we determined a cryo-EM structure of NMNAT1 bound to AMI-1 and revealed its mechanism of inhibition. This provides proof of principle for inhibiting NMNAT1 to target NAD^+^ metabolism in dependent cancers, while also highlighting that caution is warranted when interpreting studies using AMI-1 as a PRMT1 inhibitor, given its effect on NAD^+^ through NMNAT1.

**Graphical Abstract:** 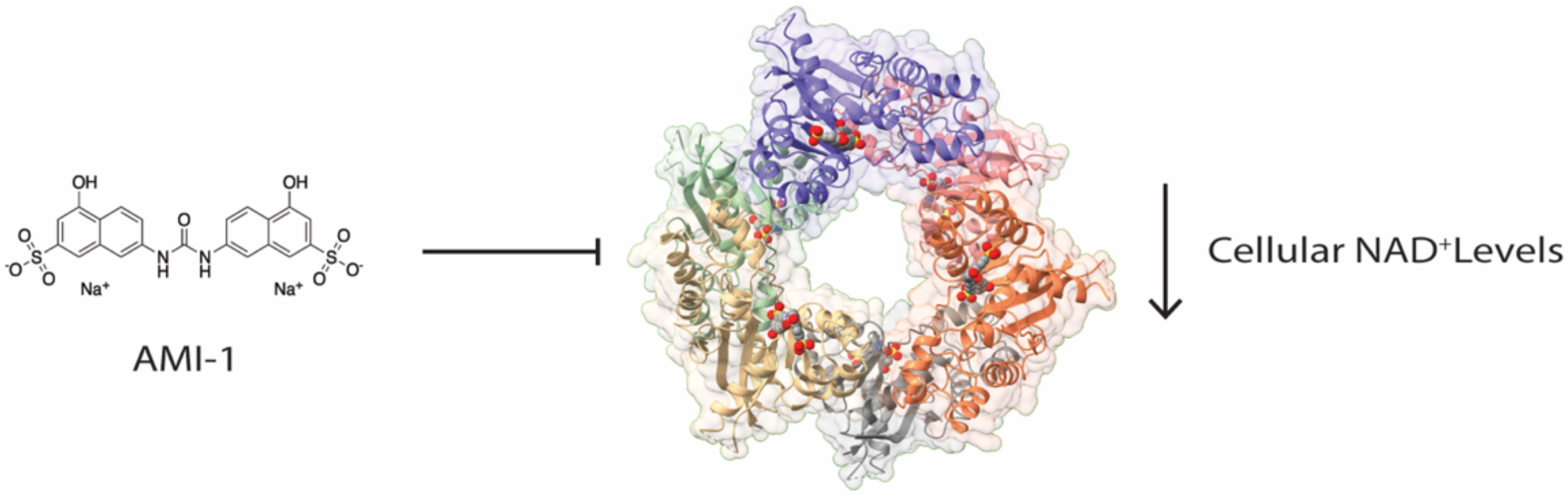

## INTRODUCTION

Nicotinamide adenine dinucleotide (NAD^+^) is a redox cofactor used by all living organisms. ^1–4^ In humans, it serves as a cofactor or substrate for metabolic enzymes, sirtuins, and PARPs ^5–7^. Nicotinamide mononucleotide adenylyltransferase (NMNAT) enzymes catalyze the synthesis of NAD^+^ from adenosine triphosphate (ATP) and nicotinamide mononucleotide (NMN)^8, 9^ (Figure 1A). Although the reaction is reversible, it is driven in the NAD^+^-generating direction, hereafter described as the forward reaction, in cells due to the high ATP concentration.^10^ There are three isoforms of NMNAT enzymes: NMNAT1, NMNAT2, and NMNAT3, localized in the nucleus, cytoplasm, and mitochondria, respectively.^11^ These isoforms are responsible for the highly compartmentalized pools of NAD^+^ in the cell and are linked to different pathologies. ^8, 12–14^

**Figure 1.**
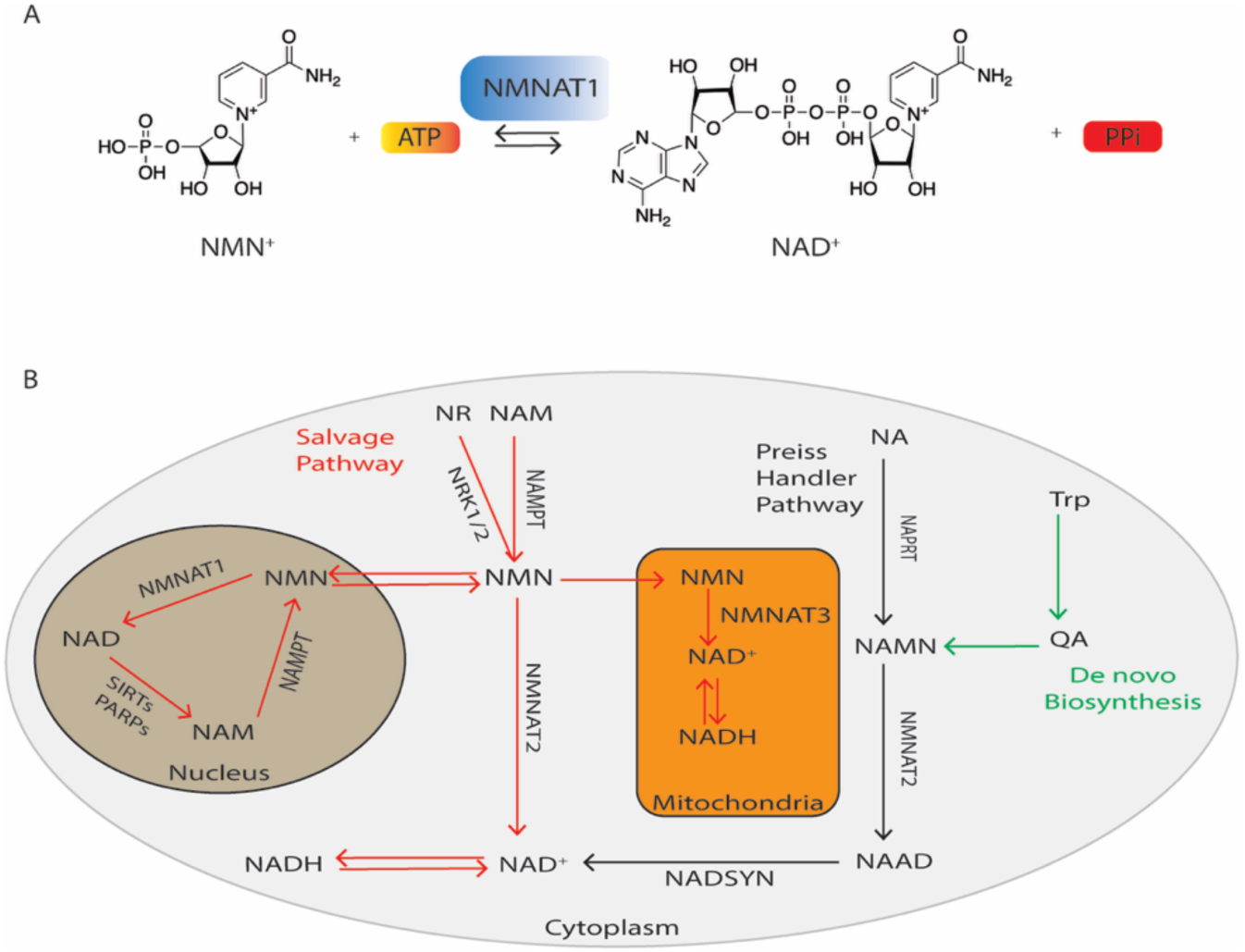
Reaction Mechanism and Cellular Compartmentalization of NMNAT1. **(A)** NMNAT1 enzymatic reaction. **(B)** NAD^+^ biosynthetic pathways and cellular compartments.

NAD^+^ is synthesized from three cellular pathways: De novo, Preiss-Handler, and the salvage pathway (Figure 1B). ^2^ The NAD^+^ salvage pathway in the nucleus is thought to be especially critical for cancer cell survival because it enables rapid proliferation by recycling NMN generated by PARP enzymes back into NAD^+^. ^15, 16^ These elevated levels of nuclear NAD^+^ produced by NMNAT1 also support DNA repair and cell survival. ^17^

Past studies have focused on targeting NAD^+^-consuming enzymes, such as sirtuins and poly (ADP-ribose) polymerases (PARPs), to regulate NAD levels.^18^ Another approach to target NAD^+^ biosynthesis is to focus on the rate-limiting enzyme in the salvage pathway, NAMPT. Potent inhibitors of NAMPT, including FK866, have entered clinical trials but were not successful due to toxicity and resistance.^19–21^ Because NAMPT is not compartmentalized, NAMPT inhibition leads to loss of NAD^+^ in all three compartments. ^22^ Recent research aims to target NMNAT enzymes for more compartment-specific NAD^+^ interventions. ^23^ A landmark study in the field showed that in AML, the salvage pathway is essential for survival, but only nuclear NAD^+^ biosynthesis is necessary. ^24^ NAMPT and NMNAT1 were found to be essential genes for AML, while NMNAT2 and NMNAT3 are not. NMNAT2 loss on the other hand has been linked to neurodegenerative diseases such as Alzheimer’s, where increased cytoplasmic NAD^+^ production from NMNAT2 has slowed disease progression. ^12^ A goal of the therapeutic strategy therefore is to target nuclear NAD^+^ levels while sparing cytoplasmic levels.

Additionally, early efforts involved developing NAD^+^ analogs such as vacor adenine dinucleotide (VAD), which inhibits NAD synthesis by targeting NMNAT enzymes. Vacor is a known metabolite that follows the same NAD^+^ biosynthesis pathway from VMN to VAD, similar to the conversion of NMN to NAD^+^. ^25^ However, as an NAD^+^ analog, it is not a selective inhibitor as it can target several enzyme classes.^26^ Another group screened 912 compounds using the reverse reaction and identified several inhibitors. ^27^ Dibromo-1,4-naphthoquinone (DBNQ) was identified as an NMNAT1 inhibitor with IC50 values of 0.76 μM and 0.26 μM in the forward and reverse reactions, respectively.^27^ However, DBNQ is a nonspecific reactive covalent compound; therefore, its cellular effect cannot be established. Currently, there are no chemical probes for NMNAT activity.

Here, we explore using biochemical and structural approaches to inhibit NMNAT1 and reduce nuclear NAD^+^ levels, as therapeutic strategy for dependent cancers. We identify Arginine Methyltransferase inhibitor-1 (AMI-1), a known Protein Arginine N-Methyltransferase 1 (PRMT1) inhibitor ^28^ as a low micromolar inhibitor of NMNAT1. We also determined a cryo-EM structure of AMI-1 bound to NMNAT1 and determined its mechanism of inhibition. Lastly, we show that AMI-1 can reduce nuclear NAD^+^ levels and can inhibit the growth of NMNAT1-dependent cancer cells.

## RESULTS

### High Throughput Screening of a Bioactive Library

To identify potential new inhibitors of NMNAT1, we overexpressed and purified human NMNAT1. We used our recombinant NMNAT1 to optimize an enzyme activity assay for the reverse reaction of NMNAT1, similar to a previously reported assay.^27^ The reverse reaction produces ATP (Figure 1A), which correlates linearly with luminescence signals from a luciferase assay. After optimizing the assay to achieve a Z’ of 0.8, we screened a known bioactives drug library (Cat# L1021) containing around 3,300 compounds. From these 3,300 compounds, we established a hit rate of ∼0.3%, resulting in 10 inhibitors that reduced NMNAT1 activity by more than 70% (Figure 2A). These compounds included natural products and other known molecules with prior biological relevance. With these 10 inhibitors, we then conducted a luciferase counter-screen to exclude those that inhibit luciferase. Seven compounds inhibited NMNAT1 but not luciferase. We repurchased these compounds and measured IC_50_ values (Figure 2B). The three most potent compounds were punicalagin, norharmane, and AMI-1 (Figure 2B, 2C Table S1). punicalagin and norharmane are both natural products. Punicalagin proved to be a highly potent compound, but we observed protein precipitation and inhibition of an unrelated enzyme.

**Figure 2.**
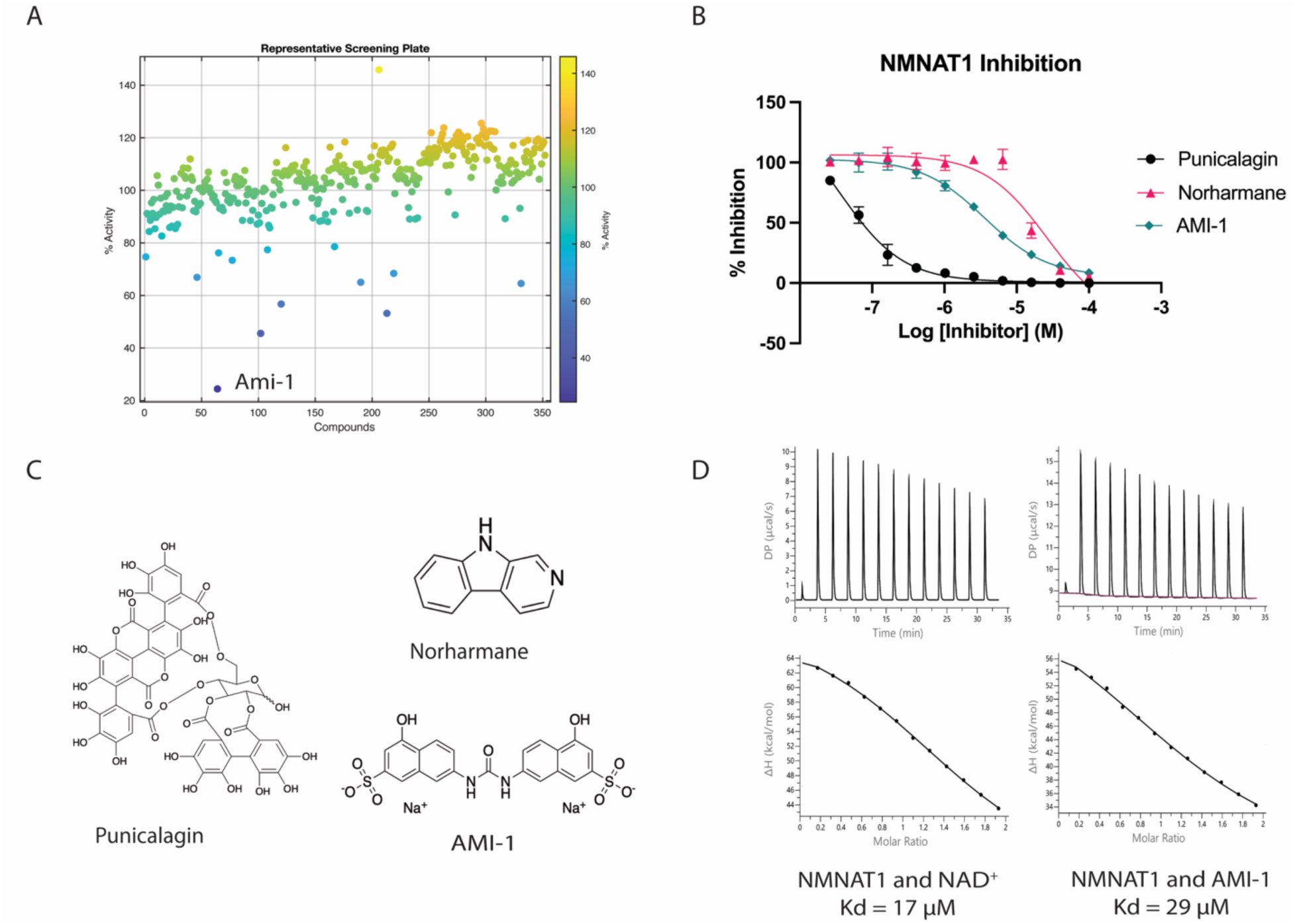
High-throughput screening and Hit Validation for NMNAT1 Inhibitors. **(A)** Representative Scatter Plot for Screening Hits from one of the screening plates. **(B)** Dose-response curves for top inhibitors from screen. Error bars show SD and n=3. **(C)** Chemical structures of top inhibitors from our screen. **(D)** ITC analysis of NAD^+^ and AMI-1 binding to NMNAT1.

Norharmane was the least potent of the hits, and its commercially available analogs showed no activity (Table S3). Therefore, we did not proceed with these compounds, as they were not viable NMNAT1 inhibitors, and focused our efforts on characterizing AMI-1.

### AMI-1 is a micromolar inhibitor of NMNAT1

AMI-1 showed micromolar inhibitory activity in both forward and reverse enzymatic reactions (Figure 2B, Table S1-S2) and demonstrated modest binding in ITC experiments (Figure 2D). The discovery of AMI-1 as an NMNAT1 inhibitor reveals a novel drug scaffold for this enzyme class. Notably, both NAD^+^ and AMI-1 exhibited endothermic binding to NMNAT1, suggesting that entropy drives these binding events (Figure 2D). We performed inhibition kinetics and found that AMI-1 is a competitive inhibitor of NMNAT1 with respect to ATP (Figure S1, Table S4).

We synthesized analogs of AMI-1 that lacked sulfonic acid moieties, with the hope of improving potency or cell permeability; however, these analogs showed negligible activity (Figure S2, Table S5), confirming the significance of the sulfonic acid for the compound’s potency.

### AMI-1 structure uncovers a novel interaction in the active site of NMNAT1

To understand how AMI-1 inhibits NMNAT1, we attempted to determine the structure of the complex. NMNAT1 has previously been crystallized with various substrates bound.^29–31^ Its structure forms a hexamer composed of two trimeric rings stacked on top of each other.

However, co-crystallization efforts of NMNAT1 and AMI-1 were unsuccessful. Therefore, we used cryo-EM to obtain the structure (Figure 3A, 3B). Initial cryo-EM experiments showed that NMNAT1 tends to adopt a preferred orientation on grids, with most particles presenting top views. To address this, we added detergent to reduce the preferred orientation and successfully obtained a 3.2 Å structure with unambiguous density for AMI-1 (Figure 3B, Table S6). We observed that the protein maintains a similar hexameric conformation to that observed in previous crystal structures, with an AMI-1 molecule at each of the six active sites (Figure 3B). Previous studies identified key catalytic residues located in the active site formed in the cleft of each monomer.^29^ AMI-1 binds inside this long channel in each monomer, occupying the space of NAD^+^ (Figure 3C). Although there is no structure of NMNAT1 bound to ATP, the adenosine monophosphate (AMP) portion of NAD^+^ overlaps significantly with AMI-1, consistent with the competitive nature of the inhibition. The only key interaction we can observe is that the sulfonic acid group of AMI-1 forms a hydrogen bond with the backbone amide glycine 156 (Figure 3D). In addition to the hydrogen bond, the sulfonic acid is positioned at the N-terminus of a helix beginning at alanine 157, whose dipole could interact favorably with the negative charge. We previously observed a similar helix-phosphate interaction in another enzyme class.^32^ On the other end of the AMI-1 symmetric dimer, the sulfonic acid points outward into the solvent. The compound induces a significant shift in the structure of the monomer by inducing a widening of the cleft. The backbone of the protein shifts by 3.3 Å (Figure 3D). We also note that there is no visible density for the final helix in NMNAT1 (residues 268-275) that is observed in the crystal structure. In the crystal structure of NMANT1 bound to NAD^+^, this helix is stabilized by an interaction with the 167-168 loop that is pushed out by AMI-1. The C-terminal helix is critical because it is not present in the otherwise highly conserved NMNAT2 isoform. Compounds that can interact with this helix might achieve selectivity over NMNAT2. Lastly, we note that modern computational programs, such as Boltz-2, failed to predict the binding site and the conformational changes induced by AMI-1, likely due to the lack of structures of similar enzymes with bound compounds and the entropic nature of the binding (Figure S4).

**Figure 3.**
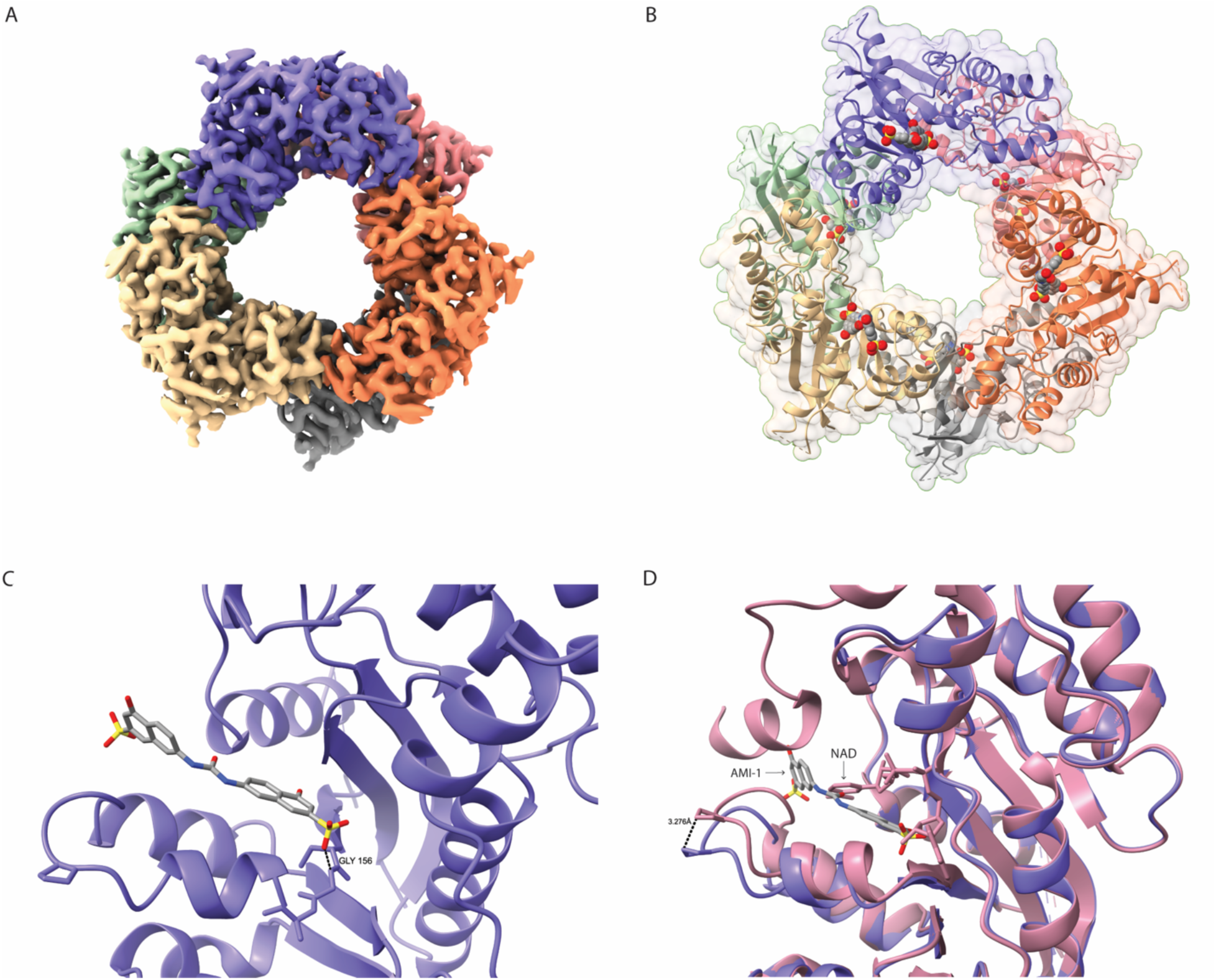
Cryo-EM Structural Insights of AMI-1 bound to NMNAT1. **(A)** Cryo-EM Electron density map of NMNAT1 with AMI-1, colored by Chain. **(B)** Surface and ribbon model of NMNAT1 with AMI-1 shown as spheres in active site **(C)** Close-up of AMI-1 interaction with glycine156 backbone in the active site of NMNAT1 **(D)** Overlay of PDB:1KQN (Pink, crystal structure of NMNAT1 bound to NAD^+^) and our structure (Blue) showing conformational change in NMNAT1 with AMI-1 bound.

### Cellular Effects of AMI-1

To further investigate the inhibitory effect of AMI-1 on NAD^+^ pools in the cell, we used a previously reporter BRET-based NAD^+^ sensors in HEK293 T-REx cells that measures nuclear, cytoplasmic, and mitochondrial NAD^+^. ^33^ Initial sensors showed toxicity, so we modified them to be doxycycline (dox)-inducible to minimize cell exposure to the sensor. We hypothesize that this might be because overexpression binds and depletes NAD^+^. We show that the known NAMPT inhibitor FK866, at 10 nM, drastically depleted NAD^+^ levels across the nucleus, cytoplasm, and mitochondria (Figure 4A). In contrast, we demonstrated that AMI-1 at 100 μM significantly reduced nuclear NAD^+^ levels while sparing the cytoplasmic and mitochondrial pools (Figure 4A). To analyze the effect of AMI-1 on the overall NAD^+^ pool in the cell and to validate our biosensor in an orthogonal assay, we utilized two cancer cell lines that depend on NMNAT1: SU-DHL-1 and NB4.^34^ Again, we established that FK866 drastically depleted NAD^+^ levels in SU-DHL-1 and NB4 cells, as well as in our normal HK-2 cells (Figure 4B). AMI-1 strongly depleted NAD^+^ levels in SU-DHL-1, while having a similar effect of NB4 and HK-2 cell lines (Figure 4B). Lastly, we assessed the viability of cancer cell lines at various concentrations of AMI-1. AMI-1 reduced cell viability more effectively in SU-DHL-1 than in NB4 or HK-2 cells (Figure 4C), which correlates with its ability to lower NAD^+^ levels in these cells.

**Figure 4.**
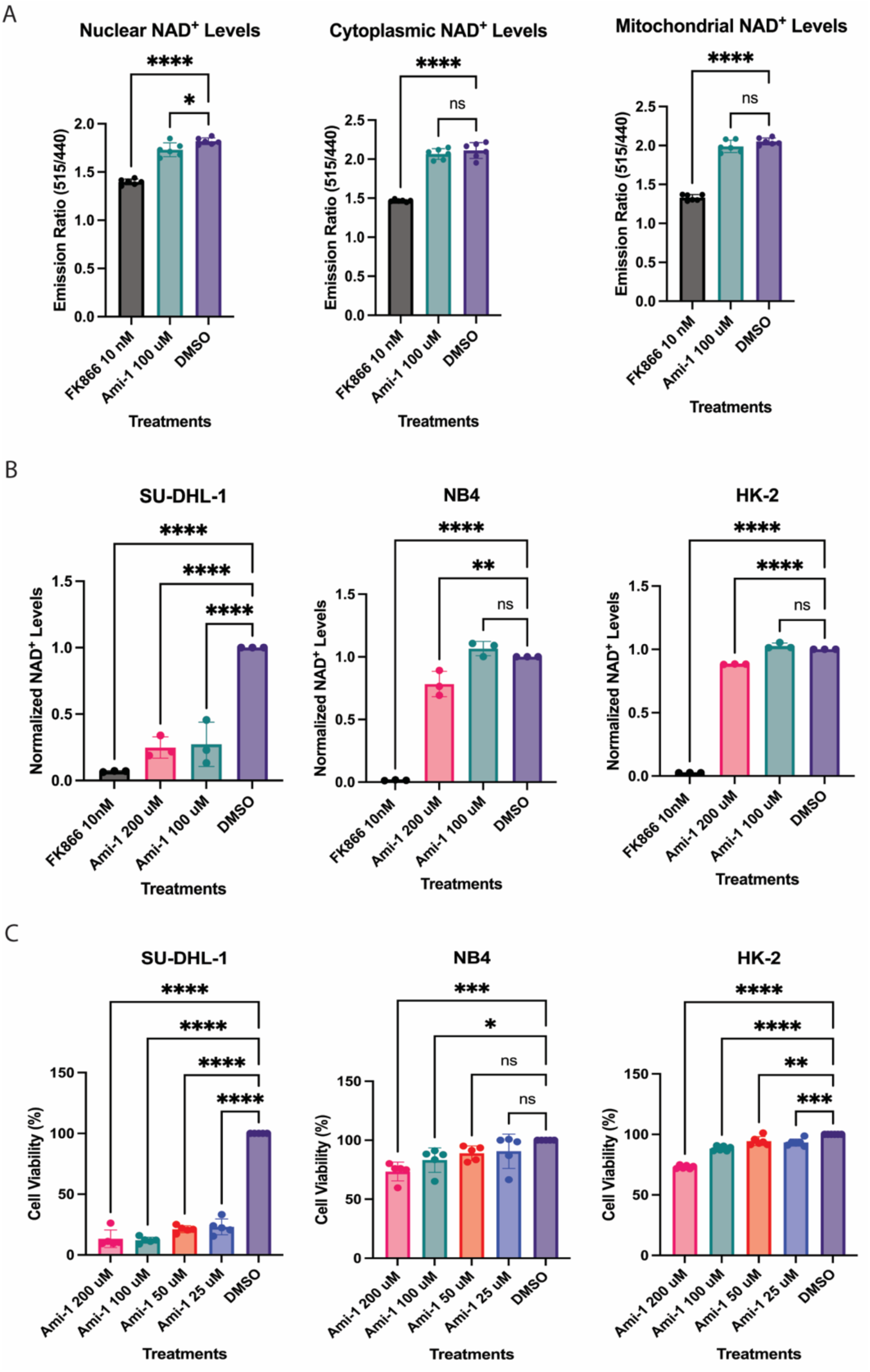
Cellular Effects of AMI-1. **(A)** Compartmentalized NAD^+^ levels using BRET-based sensor after 24-hour treatment with FK866 10nM and AMI-1 at 100 μM. Error bars show SD, n=6. **(B)** Overall NAD^+^ levels in SU-DHL-1, NB4, and HK-2 cells after 72-hour treatment with FK866 10 nM and AMI-1 at 200 μM and 100 μM from NAD glo assay. NAD^+^ levels normalized to DMSO. Error bars show SD, n=3. **(C)** Cell viability of SU-DHL-1, NB4, and HK-2 cells after 72-hour treatment with AMI-1 at a 2-fold dilution, starting at 200 μM. Error bars show SD, n=5.

## DISCUSSION

NAD^+^ biosynthesis is essential for myriad redox reactions in the cell, and NAMPT was previously identified as the rate-limiting enzyme in its production.^21^ Therefore, research focused heavily on identifying NAMPT inhibitors, but later revealed that NAD^+^ synthesis can be bypassed by precursors such as nicotinamide riboside. ^24^ Since then, NMNAT enzymes have been identified as the true gatekeepers of NAD^+^ biosynthesis and a promising target for NMNAT1-dependent cancers.^17, 24^

AMI-1 is a modestly potent, cell-permeable, and reversible compound. Its structure consists of two naphthalene rings, each substituted with a hydroxyl group and a sulfonic acid group. It inhibits NMNAT1 with an IC_50_ of 4 μM, comparable to its IC_50_ for PRMT1 of 8.8 μM.^28^

AMI-1 inhibits PRMT1 by blocking its peptide substrate binding rather than competing with the cofactor.^28, 31^ Binding studies revealed an endothermic reaction for NMNAT1. The endothermic nature of NMNAT1 may contribute to the lack of viable inhibitors, as it renders certain screening methods, such as virtual screens, more challenging. Therefore, having the first structure of NMNAT1 with a compound bound provides a good starting point for understanding how to inhibit this enzyme. Using structural and kinetic studies, we demonstrated that AMI-1 is a competitive inhibitor of ATP (Figure S1, Table S4) because it occupies the active site of NMNAT1, including both the presumptive ATP binding and the known NMN binding portions. Previous studies have determined the structure of NMNAT1 in complex with NMN, identifying key residues that stabilize the substrate and facilitate its transition states.^29^ Interestingly, it also moves away from NAD^+^, thereby occupying a portion of the pocket not observed in the NMN or NAD^+^ crystal structures, where the interior naphthalene resides before the sulfonic acid overlaps with the adenine ribose again. Our synthesized analogs demonstrate the importance of the sulfonic acid for binding, as the lack of this moiety abolished inhibitory activity (Figure S2, Table S4). This is likely due to interactions between the moiety and the glycine 156 backbone and the helix dipole. The remaining energy likely comes from entropy, which is harder to rationalize in the structure. Therefore, optimizing NMNAT1 inhibitors might be challenging, but now we have some initial insights into how to inhibit the enzyme.

FK866, a known NAMPT inhibitor, reduced overall NAD^+^ levels in the cell by approximately 90%, while AMI-1 demonstrated a modest but statistically significant reduction of nuclear NAD^+^ levels and spared cytoplasmic and mitochondrial NAD^+^ (Figure 4A). However, the differences between nuclear and cytoplasmic reductions were minimal, suggesting no significant selectivity over NMNAT2. Additionally, AMI-1 effectively depleted overall NAD^+^ levels and reduced survival in SU-DHL-1, an NMNAT1-dependent cancer cell line, showing a strong correlation between NAD^+^ levels and cell survival. (Figure 4B and 4C). However, AMI-1 was less effective at reducing viability and NAD^+^ levels in NB4 and HK-2 cells (Figure 4B and 4C). The SU-DHL-1 results show proof-of-concept for targeting NMNAT1, however, the reduced efficacy in NB4 cells and HK-2 cells confirm that a more potent compound is needed.

Future work will be needed to modify AMI-1 or identify additional scaffolds that can more potently inhibit NMNAT1 and show selectivity over other isoforms and off-targets, such as PRMT1. Currently, FK866 is a much more potent compound, so optimization of NMNAT1 inhibitors will be needed to reach that level of potency. Our attempts to investigate commercially available and designed smaller analogs of AMI-1 were unsuccessful. However, we have identified the first cellular inhibitor of NMNAT1 and show that it can reduce NAD^+^ levels in NMNAT1-dependent cells, setting the stage for the development of improved compounds.

## METHODS

### Protein expression and purification of NMNAT1

The NMNAT1 cDNA clone was obtained from DNASU. Cells were cultured in TB medium with kanamycin. A 1 L culture for protein expression was initiated with 10 mL of an overnight culture and incubated at 37°C with shaking for 2 hours, then the initial OD_600_ was measured.

OD_600_ readings were taken every 30 minutes until the OD_600_ reached 1.5. The temperature was then lowered to 16°C, and 500 µM IPTG was added to induce protein expression. Cells were harvested by centrifugation at 4°C. The pellet was resuspended and sonicated (Qsonica) in a buffer containing TBS (20 mM Tris, pH 8.0, 150 mM NaCl), PMSF (1 mM), and lysozyme (100 ug/mL), to lyse the bacteria. A column with HisPur Ni-NTA resin was washed and equilibrated with TBS containing 40 mM imidazole at pH 8.0, then washed with 20 mM Tris (pH 8.0) and 50 mM imidazole, with pH adjusted to 7.5. The solution was centrifuged and the supernatant was loaded onto the Ni-NTA resin (Thermo Scientific) for immobilized metal affinity chromatography (IMAC). NMNAT1 was eluted with 20 mM Tris (pH 8.0), 150 mM NaCl, 150 mM imidazole, and 10% glycerol. The eluate was incubated overnight with the SUMO protease and then loaded onto a Superdex 200 10/300 Increase size-exclusion column with buffer containing 25 mM HEPES pH 7.5, 500 mM NaCl, and 5 mM DTT. The protein was concentrated and stored at -80°C for assays.

### NMNAT1 Enzyme Activity Assay

To evaluate the activity of purified NMNAT1, an assay based on the reverse reaction was adapted from a previous study ^27^. This reverse reaction consumes NAD^+^ in the presence of pyrophosphate (PPi) to produce NMN and ATP. The assay was optimized with respect to buffer composition, reaction time, substrate, and protein concentration. The final buffer contained 15 mM HEPES at pH 7.5, 20 mM NaCl, 10 mM, 0.04% Triton-X, and 100 μM NAD^+^. NMNAT1 was used at a concentration of 1 nM unless otherwise noted, with an optimal reaction time of 1 hour, during which enzyme activity remained in the linear range. The substrate pyrophosphate (PPi) was at a final concentration of 100 μM in the reaction. The ATP produced is directly proportional to the enzyme activity. ATP levels were measured using the Kinase-Glo Luminescent kit (V6711) from Promega.

### High-throughput Screening of Small Molecule Inhibitors

To identify NMNAT1 inhibitors, we developed and conducted a high-throughput screen based on the enzyme activity assay described above. Our final reaction mixture consisted of 5 μL of 2x buffer and NMNAT1 (1 nM) in TBS, 4 μL of PPi, and 1 μL of compound from the APExBIO library in 10% DMSO. We used a Multidrop dispenser to transfer buffer and PPi to white Corning 384-well plates and an SPT Lab Apricot for precise pipetting of the drug plates. We performed a pilot screen with approximately 600 compounds from the bioactive library at APExBIO, followed by a full-screening assay with approximately 3,300 inhibitors. The screening assay yielded a good Z’ value and a strong assay window. From this screen, we identified 10 small-molecule inhibitors (a 0.3% hit rate) that showed greater than 70% inhibition of NMNAT1. We validated our results by performing counter-screening to ensure that small molecules did not inhibit luciferase in our assay, thereby narrowing the hit set to 7 small molecules.

### IC_50_ Reverse Reaction Validation

To evaluate the efficacy of our top small-molecule inhibitors, a concentration-response inhibition assay was used. Test inhibitors were serially diluted 2.5-fold in 10% DMSO to achieve a final starting concentration of 100 μM. The reaction was performed as outlined in the NMNAT1 enzyme activity assay protocol.

### IC_50_ Forward Reaction Validation

Drug hits were validated using the forward reaction of NMNAT1, which converts NMN and ATP into NAD^+^ and PPi. NAD^+^ production was used to optimize a resazurin assay for our counter-screen. Test inhibitors were serially diluted 2.5-fold in 10% DMSO, starting at a final concentration of 100 μM. A buffer solution was prepared with 15 mM HEPES, 20 mM NaCl, 10 mM MgCl2, 0.02% Triton-X, 100 μM NMN, 0.05% EtOH, 0.02 mg/mL diaphorase, 12.5 μM resazurin, 100 μM ATP, and 10 units of ADH in the reaction. Buffer, compounds, and 0.5 nM NMNAT1 were incubated for 30 minutes before absorbance was recorded at 570 nm.

### Isothermal Calorimetry (ITC)

The ITC protocol was adapted from previously published work.^35^ The expression and purification of NMNAT1 for isothermal titration calorimetry (ITC) experiments were performed as described above. The binding affinity of NMNAT1 for AMI-1 was assessed using a MicroCal ITC200 calorimeter (Malvern, UK). To determine the binding affinity to NMNAT1 (500 μM), 13 injections from the syringe solution were titrated into 300 μL of the cell solution (50 μM of AMI-1) with stirring at 750 rpm in a buffer containing 20 mM HEPES pH 7.4, 150 mM NaCl, 5 mM. The data were fitted using a single-binding-site model in MicroCal Origin 7.0 (Malvern).

### Electron microscopy

For electron microscopy, 3 μL of NMNAT1 full-length at a concentration of 2.5 mg/ml with 0.05% OG was applied to glow-discharged Quantifoil holey carbon grids (Cu, R1.2/1.3, 300 mesh). The grids were blotted for 2 seconds and plunged into liquid ethane using a Vitrobot plunger (4°C and 90% humidity). Cryo-EM data were collected with a Titan Krios microscope (FEI) operated at 300 kV, and images were acquired at a nominal magnification of 105,000, corresponding to a pixel size of 0.826 Å with a defocus range of −0.6 to −2 μm. The images were recorded with a Gatan K3 Direct Electron Detector in super-resolution mode at the end of a GIF-Quantum energy filter, using a slit width of 10-20 eV. A dose rate of 4.4 electrons per pixel per second and an exposure time of 6 seconds were used, yielding an accumulated dose of 50 electrons per Å².

### Image Processing and Model Building

A total of 18,166 dose-weighted micrographs of NMNAT1 were collected and imported. The processing was performed within cryoSPARC. CTF estimation for each micrograph was performed using Patch CTF estimation. From a subset of the total micrographs used for blob picking, over three million particles were selected through template picking and extracted from the micrographs. Ab initio models were generated as initial references for subsequent 3D classifications. The particles were then subjected to multiple rounds of two-dimensional (2D) and three-dimensional (3D) classification to remove poor particles; a final set of 250,000 cleaned particles was used for NU-refinement with D3 symmetry, yielding a final map at 3.23 Å resolution. Structures were built in Coot based on previous crystal structures of full-length NMNAT1 ^36^, and further refined using PHENIX. UCSF Chimera ^37^ was used to generate the figures shown in the manuscript.

### Cryo-EM refinement

Using our cryo-EM structure, we fitted a model with Phenix ^38^ dock_in_map and refined it iteratively through manual adjustments in Coot ^39^ and Phenix real_space_refine, including B-factor refinement and rotamer correction.

### Stable Cell Line Generation Cloning

Ratiometric NAD^+^ sensor vectors were used as PCR templates to amplify Nuclear-Olive, Cytoplasmic-Olive, and Mitochondrial-Olive cDNA sequences. ^33^ These were cloned into the pcDNA5-FRT-TO-GFP vector ^40^, replacing the GFP gene, using Gibson ligation with the appropriate primers for each gene and the vector.

### Primer sequences

The pcDNA-FRT-TO-GFP vector primers were: GGTGGCAAGCTTAAGTTTAAACGCTAGAGTCCGG and 5’-GGATCCACTAGTCCAGTGTGGTGGAATTCTGC. The nuclear, cytoplasmic, and mitochondrial olive N-terminal primers were, respectively: 5’-CCGGACTCTAGCGTTTAAACTTAAGCTTGCCACC atggatccaaaaaagaagagaaaggtagcctccctg, 5’-CCGGACTCTAGCGTTTAAACTTAAGCTTGCCACC atgctgcagaatgaactggcactgaagctc, and 5’-CCGGACTCTAGCGTTTAAACTTAAGCTTGCCACC atgtctgttctgactcctctgctgctccg. The olive C-terminal primer was: 5’-GCAGAATTCCACCACACTGGACTAGTGGATCC tcactcatacgggatgatcacatgaatatcgattttcaggc.

### Stable Cell Lines

The Flp-In™ 293 T-REx cells (Thermo Fisher, R78007) were cultured at 37°C with 5% CO₂ in DMEM supplemented with 10% FBS that is free of tetracycline. Olive/pcDNA5-FRT-TO vectors were transfected into the Flp-In™ 293 T-REx cells cultured in 6-well plates, using 0.2 μg of Olive vector and 1.8 μg of Flippase vector, pOG44 (ThermoFisher, V600520), at a 1:9 ratio, with 6 μL of PEI (1 mg/mL, pH 6.7). After 16 hours, the cell culture medium was replaced with fresh medium. On day 3, one-quarter of the cells from each well were transferred to a 15 cm dish and grown overnight. Stable cell lines were selected by culturing in medium containing 150 μg/mL of hygromycin B Gold (InvivoGen, #ant-hg-1). Surviving colonies were then trypsinized from the dish using 8×8 mm Pyrex cloning cylinders (Corning, 31668) and screened for Nanoluciferase activity.

### Nano-BRET Cell Assay

The developed stable cell lines included nuclear, cytoplasmic, and mitochondrial NAD^+^ sensors. These sensors were used to measure NAD^+^ pools in the nucleus, cytoplasm, and mitochondria. Three 96-well plates (Corning) were seeded with 0.01 × 10^6^ cells per well using tetracycline-free DMEM, supplemented with doxycycline (1:1000) to induce expression of the NAD^+^ sensors.

The cells were treated with FK866 (10 nM) and AMI-1 (100 μM) for 24 hours. The tissue culture plates were equilibrated to room temperature, and Promega Nano-Glo (N1110) luciferase reagent was diluted 1:1000 with HBSS and added to the wells at a 1:1 ratio. The reaction plates were mixed and incubated for 3 minutes, after which bioluminescence was measured using a Nivo microplate reader, with NLuc emission detected at 410 nm and mScarlet-I at 515 nm.

### NAD-glo Assay

As an orthogonal assay, the Promega NAD/NADH-Glo assay (G9071) was used to assess the effect of AMI-1 on NAD^+^ levels in different NMNAT1-dependent cell lines. The SU-DHL-1 and NB4 cancer cell lines were used as disease models and HK-2 cells as the healthy control. White 96-well plates (Corning) were seeded with 0.003 × 10^6^ cells in 100 μL. Wells were treated with FK866 at 10 nM and AMI-1 at 200 μM and 100 μM after 24 hours. The cells were incubated with the treatments for 72 hours. Medium was removed, and the cells were resuspended in 50 μL PBS, then lysed with 50 μL base solution containing 1% DTAB. To measure NAD+, 25 μL of 0.4 N HCl was added and heated to 60°C for 15 minutes, then cooled to room temperature, followed by 25 μL of 0.5 M Tris base. NAD/NADH-Glo reagent was prepared and added to the treatment wells at a 1:1 ratio to detect NAD^+^ levels. Luminescence was recorded using a Nivo microplate reader after a 30-minute incubation.

### WST-8 Assay

The Cell Counting Kit-8 (CCK-8, AbCam) was used to assess cell viability of the NMNAT1-dependent cancer cell lines SU-DHL-1 and NB4 in the presence of AMI-1. White 96-well plates (Corning) were seeded with 0.003 × 10^6^ cells in 100 μL and treated with various concentrations of AMI-1 (200 μM, 100 μM, 50 μM, and 25 μM) for 72 hours. After treatment, 10 μL of CCK-8 solution was added to each well and incubated for 1 hour before reading. The plate was mixed gently, and the absorbance was measured at 450 nm using a Victor Nivo microplate reader.

## Acknowledgements

Research reported in this publication was supported by the National Institute of General Medical Sciences of the National Institutes of Health under Award Numbers R35GM124838 (to M.B.L.). Some of this work was performed at the National Center for CryoEM Access and Training (NCCAT) and the Simons Electron Microscopy Center located at the New York Structural Biology Center, supported by the NIH Common Fund Transformative High Resolution Cryo-Electron Microscopy program (U24 GM129539, and NIGMS R24 GM154192) and by grants from the Simons Foundation (SF349247) and NY State Assembly. Some of this work was performed at the Simons Electron Microscopy Center at the New York Structural Biology Center, with major support from the Simons Foundation (SF349247). This work was supported in part by the computational and data resources and staff expertise provided by Scientific Computing and Data at the Icahn School of Medicine at Mount Sinai, and by the Clinical and Translational Science Award (CTSA) grant UL1TR004419 from the National Center for Advancing Translational Sciences. Molecular graphics and analyses performed with UCSF ChimeraX, developed by the Resource for Biocomputing, Visualization, and Informatics at the University of California, San Francisco, with support from National Institutes of Health R01-GM129325 and the Office of Cyber Infrastructure and Computational Biology, National Institute of Allergy and Infectious Diseases.

## Supporting Information

Includes Supplementary Tables and Figures.

