## Supplemental Data for "Structural and biochemical characterization of a novel inhibitor of NMNAT1, the gatekeeper of nuclear NAD^+^ biosynthesis"

**Supporting Information**

Carisse Lansiquot,[1#] Ruoxi Wu,[1#] Joanna P. Davies,[1] Xiangyang Song,[1,2,3] H Ümit Kaniskan,[1,2,3] Jian Jin,[1,2,3] Michael B. Lazarus*[1,2]

[1]Department of Pharmacological Sciences, Icahn School of Medicine at Mount Sinai, New York, New York 10029
[2]Mount Sinai Center for Therapeutics Discovery, Icahn School of Medicine at Mount Sinai, New York, New York 10029
[3]Departments of Oncological Science and Neuroscience, The Mount Sinai Tisch Cancer Center, Icahn School of Medicine at Mount Sinai, New York, NY 10029

**Contents:**
**Supplementary Tables**
**Supplementary Figures**
**Supplementary Methods**
**References**

**Table S1. Reverse $IC_{50}$ Values of NMNAT1 inhibitors**

| Compound Name | Structure | IC50 Value [μM] | 95% CI [μM] |
|---|---|---|---|
| Punicalagin | 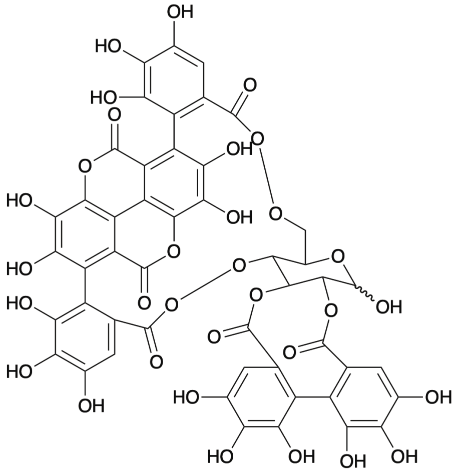 | 0.36 | 0.02 - 0.05 |
| Norharmane | 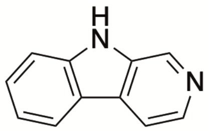 | 26 | 14.5 – 51.8 |
| AMI-1 | 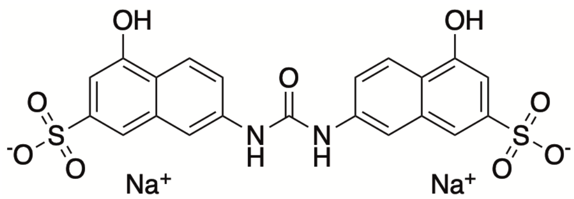 | 4 | 2.93 – 4.4 |

**Table S2. Forward $IC_{50}$ Value of AMI-1 against NMNAT1.**

| Compound Name | Structure | IC50 Value [µM] | 95% CI [µM] |
|---|---|---|---|
| AMI-1 | 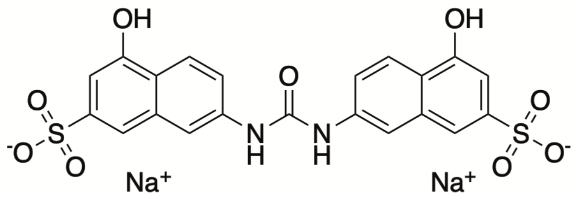 | 5 | 4.44 – 7.81 |

**Table S3. IC$_{50}$ Values of Norharmane Analogs against NMNAT1.**

| Compound Name | Structure | IC50 Value [µM] | 95% CI [µM] |
|---|---|---|---|
| Norharmane | 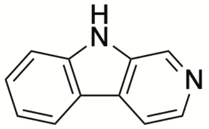 | 26 | 5.99 - 16 |
| Harmol | 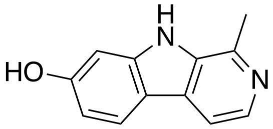 | >100 | |
| Harmane | 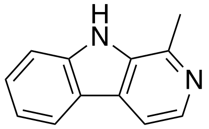 | >100 | |
| Harmine | 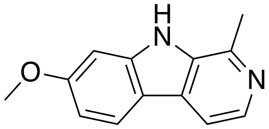 | >100 | |
| 3-bromo-9H-pyrido[2,3-b]indole | 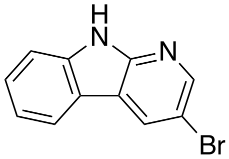 | >100 | |
| 5H-pyrido[4,3-b]indole | 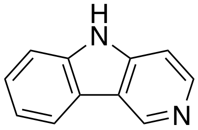 | >100 | |

**Table S4. Inhibition kinetics of AMI1 against NMNAT1 with respect to ATP.** Calculated kinetics constants from nonlinear regression are shown.

| AMI-1 concentration [μM] | Vmax | 95% CI [μM] | Km | 95% CI [μM] |
|---|---|---|---|---|
| 0 | 8.0 | 7.66 – 8.44 | 29.93 | 22.91 – 38.49 |
| 1.85 | 6.9 | 5.92 – 8.00 | 36.56 | 15.74 – 76.30 |
| 5.5 | 5.8 | 5.09 – 6.64 | 48.51 | 24.93 – 87.79 |
| 16.6 | 6.2 | 5.37 – 7.30 | 169.4 | 97.18 – 293.1 |

**Table S5. $IC_{50}$ Values of AMI-1 Analogs against NMNAT1.**

| Compound Name | Structure | IC50 Value [µM] | 95% CI [µM] |
|---|---|---|---|
| AMI-1 | 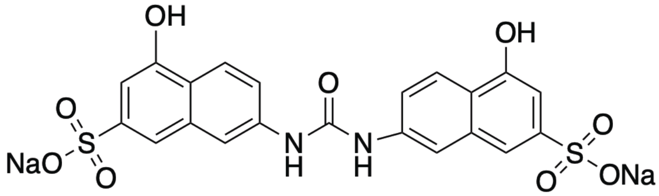 | 4 | 0.13 – 0.17 |
| XS297-25 | 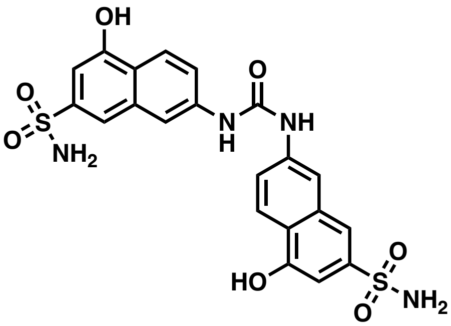 | >100 | |
| XS297-26 | 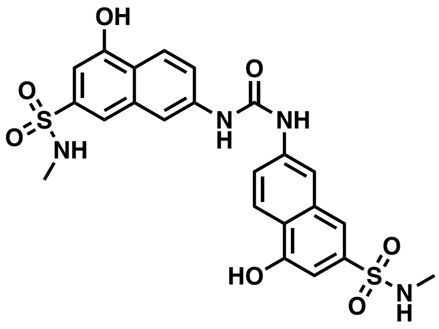 | >100 | |

**Table S6. Cryo-EM data collection, refinement, and validation statistics**

| | NMNAT1/AMI-1 |
|---|---|
| **Data collection and processing** | |
| Magnification | 105,000 |
| Voltage (kV) | 300 |
| Electron exposure (e–/Å$^2$) | 50.30 |
| Defocus range (μm) | -0.6 to -2.0 |
| Pixel size (Å) | 0.826 |
| Symmetry imposed | D3 |
| Initial particle images (no.) | 3,268,846 |
| Final particle images (no.) | 250,000 |
| Map resolution (Å) | 3.23 |
| FSC threshold | 0.143 |
| Map resolution range (Å) | 0-8.55 |
| **Refinement** | |
| Initial model used (PDB code) | |
| Model resolution (Å) | 3.23 |
| FSC threshold | 0.143 |
| Map sharpening *B* factor (Å$^2$) | -205.5 |
| Model composition | |
| Non-hydrogen atoms 10686 | |
| Protein residues | |
| Nucleotides | 1296 |
| | 0 |
| *B* factors (Å$^2$) | |
| Protein | 19.98/125.74/59.45 |
| Nucleotides | 0 |
| R.m.s. deviations | |
| Bond lengths (Å) | 0.002 |
| Bond angles (°) | 0.375 |
| Validation | |
| MolProbity score | 1.07 |
| Clashscore | 2.20 |
| Poor rotamers (%) | |
| Ramachandran plot | |
| Favored (%) | 97.64 |
| Allowed (%) | 2.36 |
| Disallowed (%) | 0 |

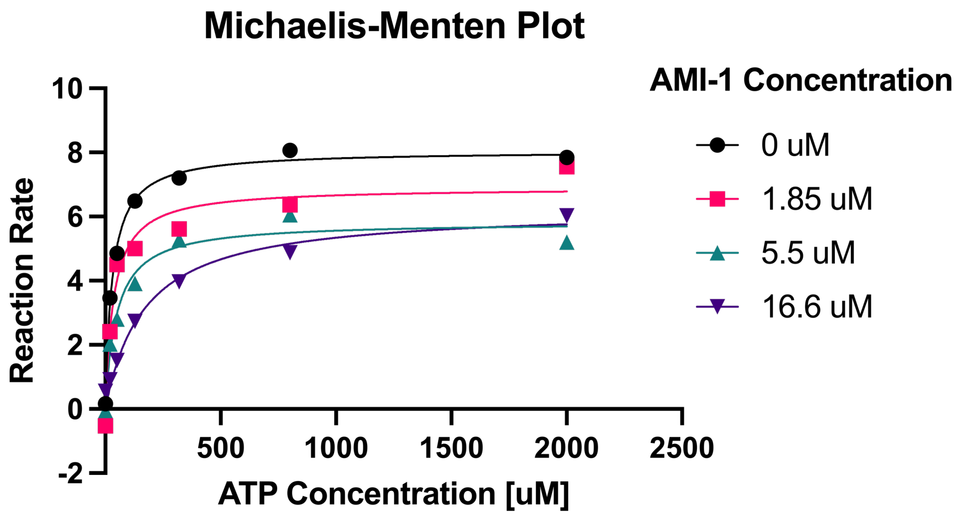


**Figure S1. Michaelis-Menten inhibition kinetics of AMI-1 against NMNAT1.** Reaction Kinetics of AMI-1 at various concentrations compared to ATP, n=1.

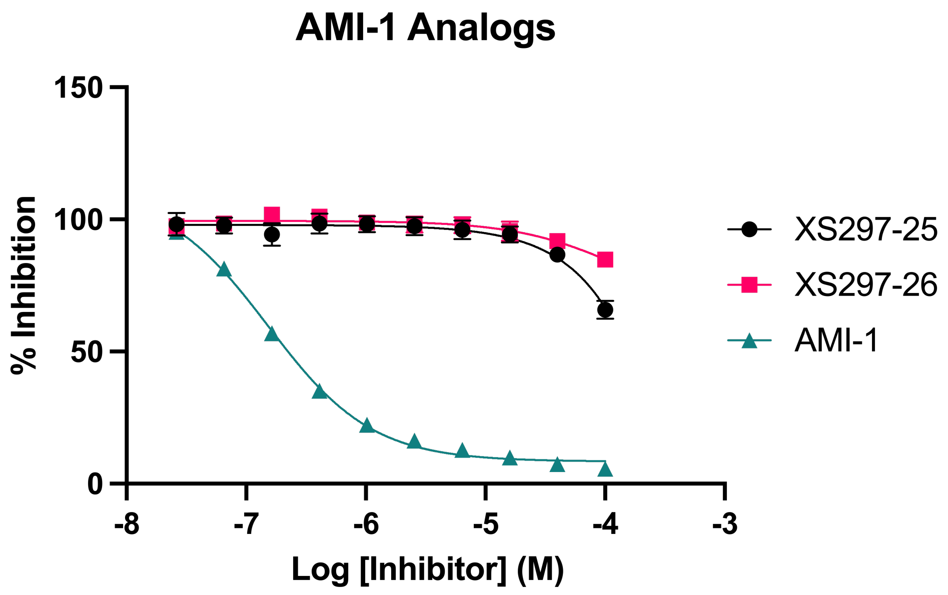


**Figure S2. $IC_{50}$ Plots for synthesized AMI-1 analogs against NMNAT1.** Dose response curve for AMI-1 synthesized analogs starting at 100 µM with 2.5-fold dilution. Error bars show SD, n=3.

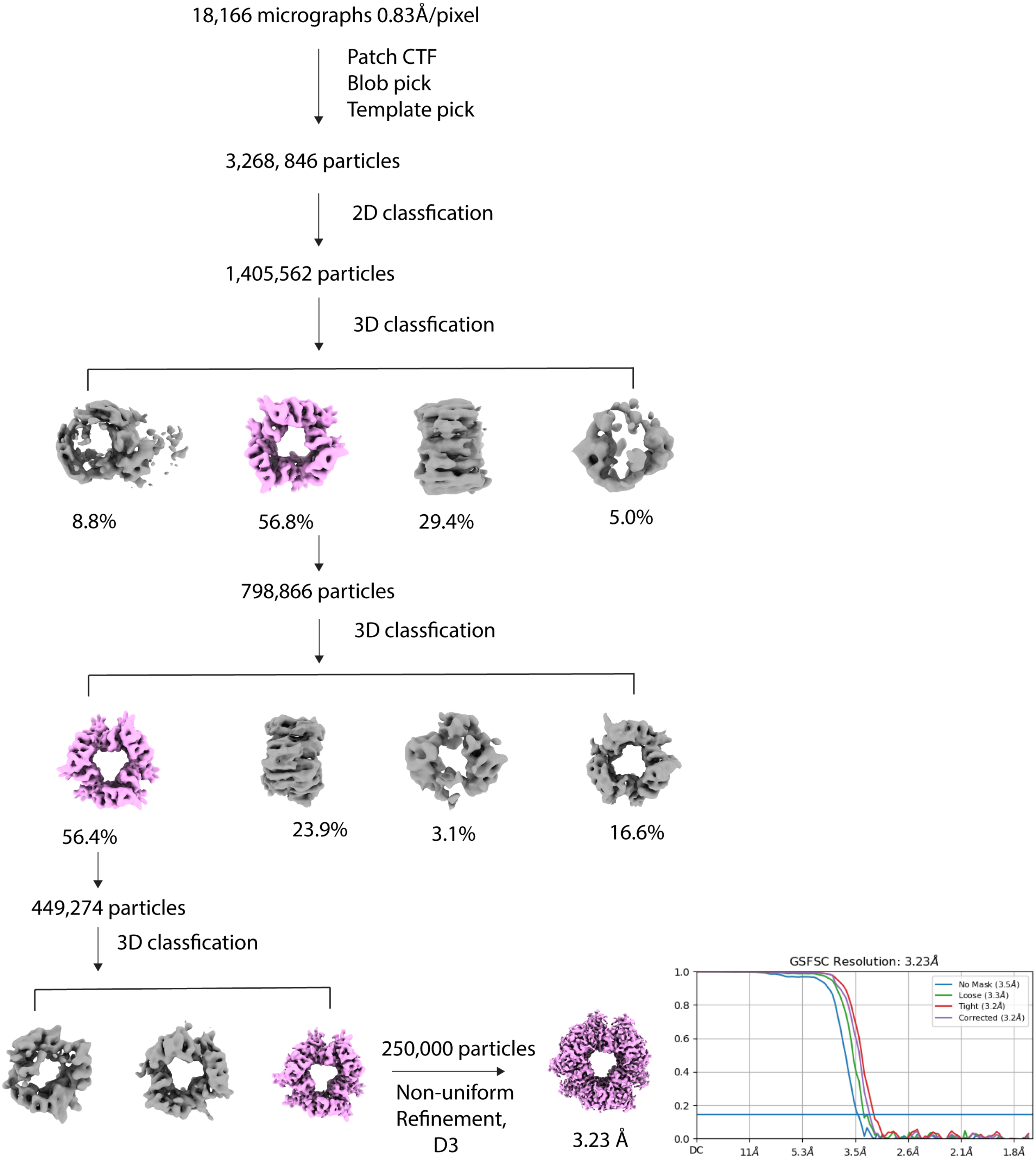


**Figure S3. Flow Chart of Sorting and Classifications used to process NMNAT1-AMI-1 Complex.**

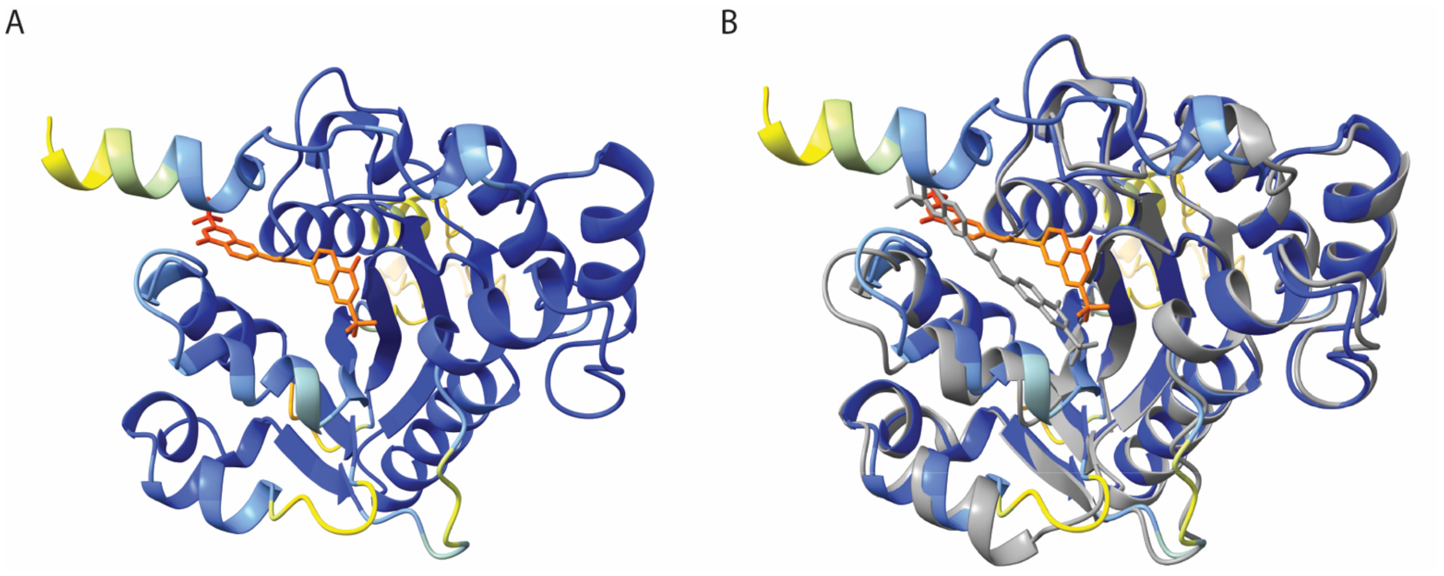


**Figure S4. (A)** Boltz-2 Model Prediction of AMI-1 bound to the active site of NMNAT1. Compound is shown as sticks in orange. **(B)** Overlay of Boltz-2 Model and our structure. Our structure is shown in grey, with AMI-1 shown as sticks.

**Synthesis of AMI-1 Analogs**

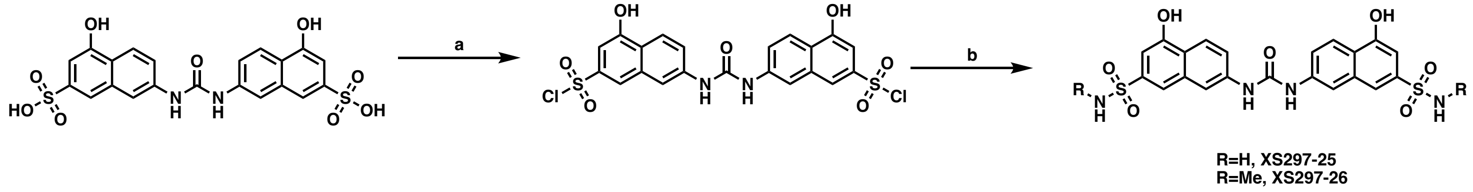


**Scheme 1(a)** $POCl_3$, 50 °C; (b) $RNH_2$, DIPEA, DMAP.

To 542 mg (1.0 mmol) of 7,7'-(carbonylbis(azanediyl))bis(4-hydroxynaphthalene-2-sulfonic acid) suspended in 5 mL of phosphorus oxychloride. The stirred suspension was heated at 50 °C, 12 h. Then the cloudy suspension was poured onto 200 mL of ice and this suspension placed in a second ice bath. After the ice had melted, the precipitate was collected by vacuum filtration and was washed with water. Drying provided 7,7'-(carbonylbis(azanediyl))bis(4-hydroxynaphthalene-2-sulfonyl chloride) as a brown solid. To a solution of the 7,7'-(carbonylbis(azanediyl))bis(4-hydroxynaphthalene-2-sulfonyl chloride) (10.8 mg, 0.02 mmol) and $RNH_2$ (70% in MeOH) in MeOH(1 mL) was added DIPEA (10.3 mg, 0.08 mmol) and DMAP (2.4 mg, 0.02 mmol). The mixture was stirred at room temperature for 3 h followed by purified by prep-HPLC to yield final compound.

**7,7'-(carbonylbis(azanediyl))bis(4-hydroxynaphthalene-2-sulfonamide)** (XS297-25) Brown powder (2.1 mg, 20% yield)$^1$H NMR (400 MHz, Methanol-$d_4$) δ 8.11 (d, $J$ = 9.1 Hz, 2H), 8.09 – 8.04 (m, 2H), 7.74 (s, 2H), 7.49 (d, $J$ = 2.0 Hz, 1H), 7.47 (d, $J$ = 2.1 Hz, 1H), 7.04 (d, $J$ = 1.6 Hz, 2H). HRMS (ESI-TOF) m/z: $[M+H]^+$ calcd for $C_{21}H_{19}N_4O_7S_2$, 503.0690; found: 503.0697.

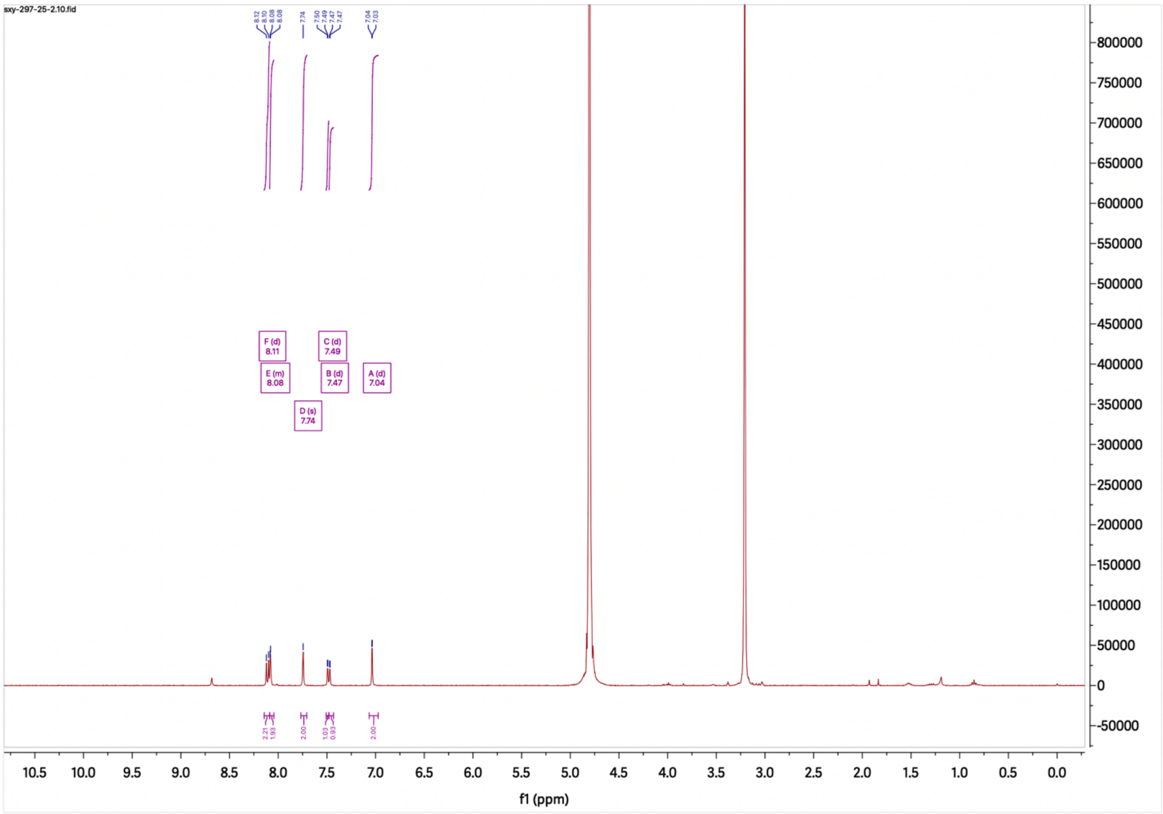


[1]H NMR Spectrum of **XS297-25**

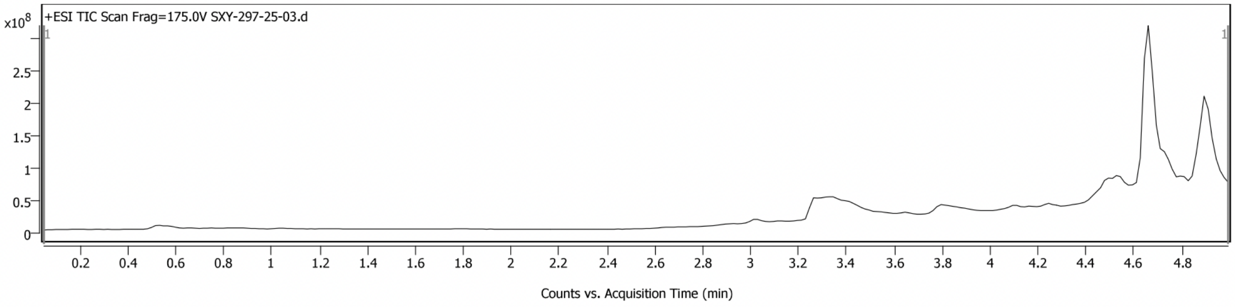


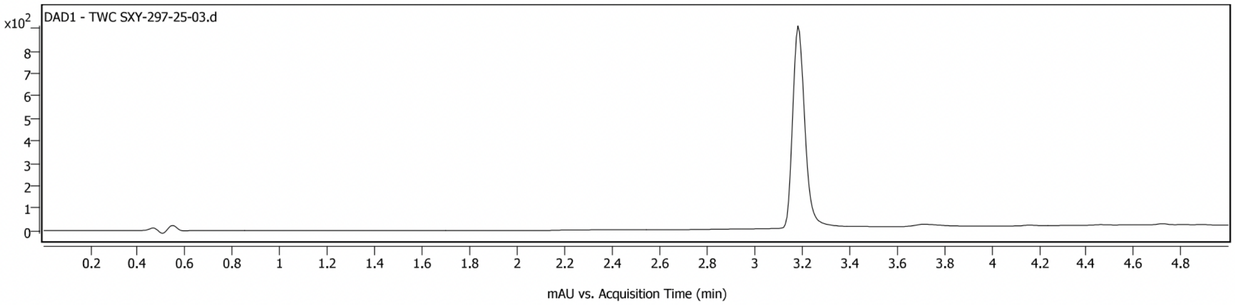


**Sample Spectra**

**+ Scan (rt: 3.278 min)**

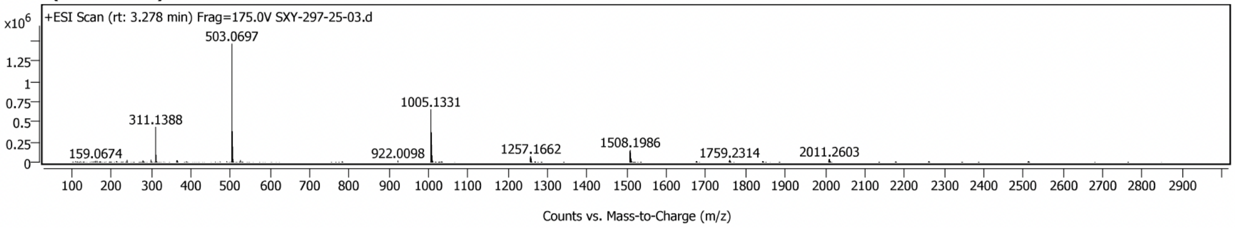


LC-MS Spectrum of **XS297-25**

**7,7'-(carbonylbis(azanediyl))bis(4-hydroxy-*N*-methylnaphthalene-2-sulfonamide)** (XS297-26), Yellow powder (5.8 mg, 55% yield). $^1$H NMR (400 MHz, Methanol-$d_4$) δ 8.18 (d, $J$ = 9.1 Hz, 2H), 8.15 (d, $J$ = 2.1 Hz, 2H), 7.74 (s, 2H), 7.58 (d, $J$ = 2.1 Hz, 1H), 7.56 (d, $J$ = 2.1 Hz, 1H), 7.02 (d, $J$ = 1.6 Hz, 2H), 2.54 (s, 6H). HRMS (ESI-TOF) m/z: $[M+H]^+$ calcd for $C_{23}H_{13}N_4O_7S_2$, 531.1003; found: 531.1004.

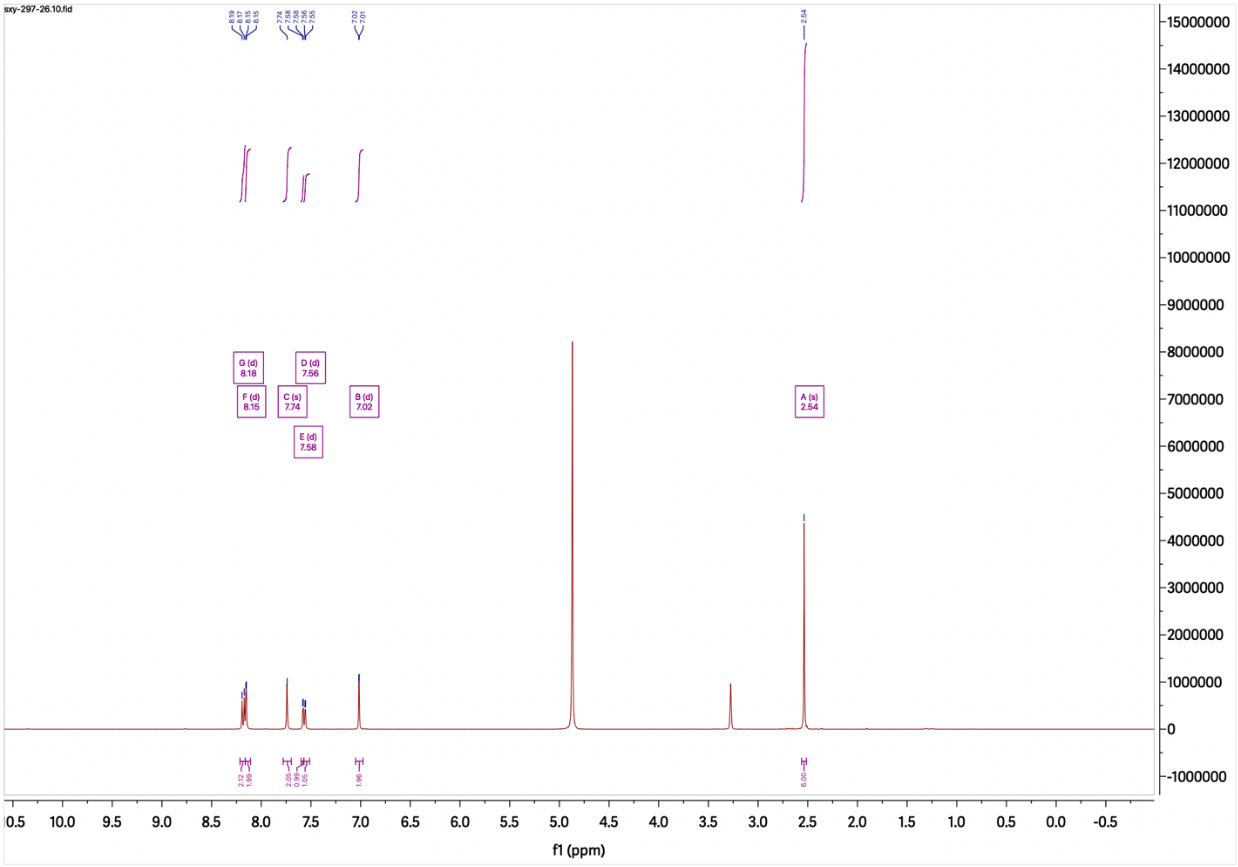


$^1$H NMR Spectrum of **XS297-26**

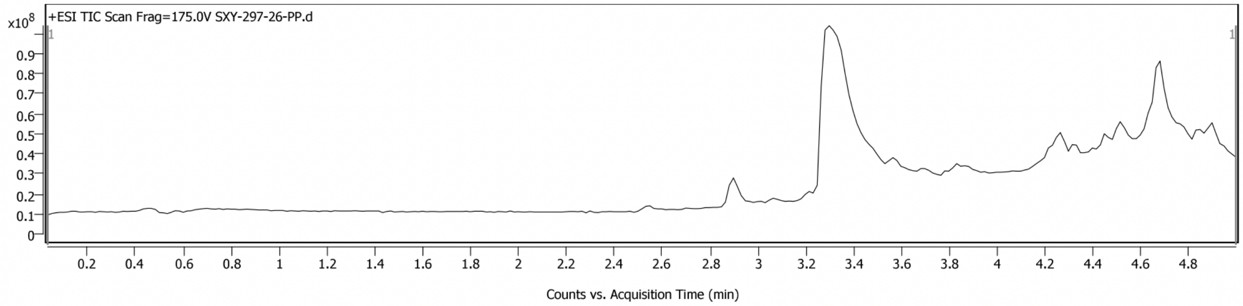


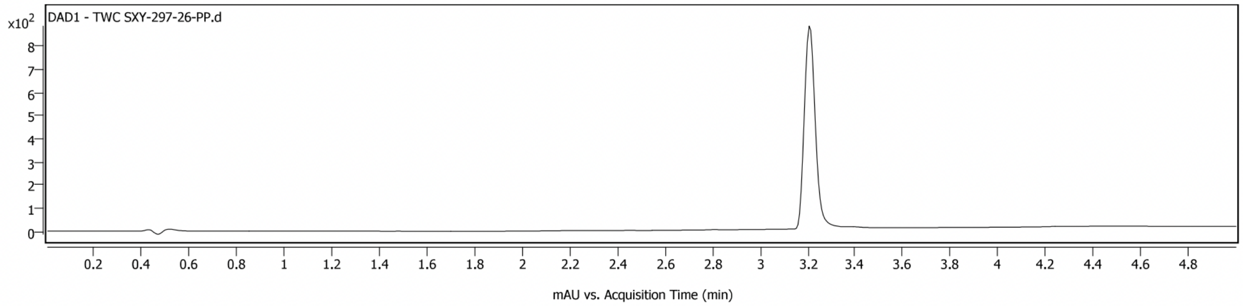


**Sample Spectra**

**+ Scan (rt: 3.281 min)**

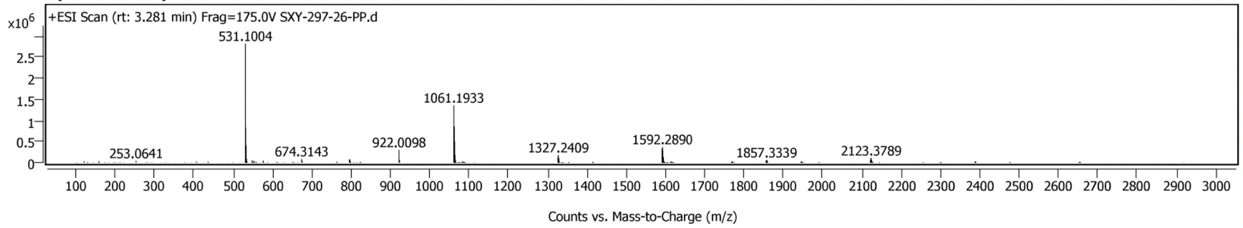


LC-MS Spectrum of **XS297-26**
